# Rapid Identification of Stable Clusters in Bacterial Populations Using the Adjusted Wallace Coefficient

**DOI:** 10.1101/299347

**Authors:** Dillon OR Barker, João A Carriço, Peter Kruczkiewicz, Federica Palma, Mirko Rossi, Eduardo N Taboada

## Abstract

Whole-genome sequencing (WGS) of microbial pathogens has become an essential part of modern epidemiological investigations. Although WGS data can be analyzed using a number of different approaches, such as traditional phylogenetic methods, a critical requirement for global systems for pathogen surveillance is the development of approaches for transforming sequence data into WGS-based subtypes, which creates a nomenclature that describes their higher-order relationships to one another. To this end, subtype similarity thresholds are needed to define clusters of subtypes representing lineages of interest. WGS-based subtyping presents a challenge since both the addition of novel genome sequences and small adjustments in similarity thresholds can have a dramatic impact on cluster composition and stability. We present the Neighbourhood Adjusted Wallace Coefficient (nAWC), a method for evaluating cluster stability based on computing cluster concordance between neighbouring similarity thresholds. The nAWC can be used to identify areas in in which distance thresholds produce robust clusters. Using datasets from *Salmonella enterica* and *Campylobacter jejuni*, representing strongly and weakly clonal bacterial species respectively, we show that clusters generated using such thresholds are both stable and reflect basic units in their overall population structure. Our results suggest that the nAWC could be useful for defining robust clusters compatible with nomenclatures for global WGS-based surveillance networks, which require stable clusters to be defined that both harness the discriminatory power of WGS data while allowing for long-term tracking of strains of interest.

## Introduction

Molecular subtyping of microbial pathogens has become integral to modern epidemiological investigations of infectious disease, playing a key role in public health efforts worldwide (Foxman and Riley, 2001). Subtyping is used to discriminate isolates from the same species and to identify clusters of similar isolates based on their genetic relatedness. In the context of molecular surveillance networks such as PulseNet, this information has made it possible to confirm possible disease outbreaks, to identify outbreak cases that may not have been possible to distinguish among a background of sporadic cases, and to further link cases to potential sources of infection even in the absence of clear epidemiological linkage (Boxrud et al., 2010; Tauxe et al., 2010).

Over the last several decades, pathogen subtyping approaches have evolved from methods based on assessing phenotypic characteristics to methods based on genetic variation detectable through gel electrophoresis and sequencing. More recently, advances in a new generation of high-throughput sequencing methods has led to the increasing adoption of whole-genome sequencing (WGS) as a primary method for the characterization of microbial pathogens (Gilmour et al., 2013; Köser et al., 2012). As traditional subtyping methods give way to whole-genome-based analyses through large-scale adoption of WGS worldwide, there is an increasing recognition that genomics-based global surveillance systems for pathogen tracking are becoming both feasible and essential (Struelens and Brisse, 2013).

Although WGS data can be analyzed using a number of different approaches, including traditional phylogenetic methods, a critical requirement for global systems for pathogen surveillance is the development of approaches for transforming sequence data into WGS-based subtyping data. The process of developing a WGS-based subtyping schema requires a standardized methodological approach for defining subtypes and a nomenclature for describing their higher-order relationships to one another. Together, these allow the rapid analysis of data, a simplified means of contextualizing results, and >an efficient process for exchanging this information, facilitating a rapid response aimed at disease prevention and control (Taboada et al., 2017).

The ‘gene-by-gene’ approach, an extension of the original Multi-Locus Sequence Typing (MLST) approach of Maiden et al. (Maiden et al., 1998) when applied to WGS data, is rapidly emerging as one of the leading approaches proposed for global surveillance using WGS-based subtyping data (Sheppard et al., 2012). In the context of WGS analysis, this increasing acceptance is due to several features that made traditional MLST one of the most widely adopted subtyping methods for investigating the population structure and epidemiology of a wide range of bacteria (Maiden, 2006). Because MLST indexes sequence variation such that each locus – a gene or gene fragment – is used as the basic unit of comparison, this approach can be applied to the development of highly standardized analytical workflows (Sheppard et al., 2012). At the same time, the MLST approach can be used to develop hierarchical subtyping schemes suitable for studying strain relationships at a range of phylogenetic depths by adjusting the number and type of loci interrogated (Maiden et al., 2013). From its inception, the MLST approach was designed to generate subtyping data from sequence information in the context of a clear and consistent nomenclature rooted on population genetics principles. The importance of this feature cannot be underestimated, as nomenclatures used to describe subtyping schemes are essential to interlaboratory information sharing, a critical feature for global WGS-based surveillance (Nadon et al., 2017).

An important consideration in the application of any subtyping scheme is the process for defining subtyping clusters composed of strains likely to be related, which is generally achieved through the application of a distance or similarity threshold for cluster definition (Tenover et al., 1995). Optimization of this parameter has been a perennial challenge in the field of molecular epidemiology, impacting both outbreak investigations, which require clusters to be defined for outbreak inclusion/exclusion, and surveillance, which requires clusters to be defined for longitudinal tracking of strains of interest. WGS-based subtyping presents a challenge since both the addition of novel genome sequences and small adjustments in similarity thresholds can have a dramatic impact on cluster composition and stability.

We present the Neighbourhood Adjusted Wallace Coefficient (nAWC), a method for evaluating cluster stability based on computing partitioning concordance between clusters derived from neighbouring distance thresholds. We show, using datasets from *Salmonella enterica* and *Campylobacter jejuni*, that this approach can be used to identify robust clusters that can form the basis for stable nomenclatures, an essential component of global surveillance networks for pathogen tracking.

## Methods

### *Salmonella enterica* and *Campylobacter jejuni* genome sequences

For *Salmonella enterica*, henceforth referred to as *Salmonella*, WGS data (n=91,295) was obtained from several sources. These included: genome assemblies from EnteroBase (n=86,913); genome assemblies previously described in (Yachison et al., 2017; Yoshida et al., 2016) (n=3,795); raw reads from the Short Read Archive (SRA; http://www.ncbi.nlm.nih.gov/sra) at the National Center for Biotechnology Information (n=3,818), which were assembled using SPAdes (v3.11) (Bankevich et al., 2012). For *Campylobacter jejuni*, henceforth referred to as *Campylobacter*, raw WGS reads for genomes used in this study (n=6,784) were obtained from the SRA. Draft genome assemblies were produced using the INNUca pipeline (https://github.com/b-ummi/innuca). Genomes were selected to represent as much of diversity in genetics and provenance as currently available.

### Core Genome Multi-Locus Sequence Typing (cgMLST)

For genome-scale MLST analysis we used a method derived from the approach for whole genome MLST (wgMLST) previously described by Sheppard et al. (Sheppard et al., 2012) but focusing on core genes, which has been termed ‘core genome MLST’ (cgMLST). For *Salmonella* we used a 330 locus cgMLST schema previously described and implemented in the *Salmonella In Silico* Typing Resource (SISTR) (Yoshida et al., 2016) using the command-line version of the platform (sistr_cmd v1.0.2; https://github.com/peterk87/sistr_cmd). For *Campylobacter*, we developed a prototype cgMLST scheme as described below and cgMLST analysis was performed using the Microbial *In Silico* Typing (MIST) platform (Kruczkiewicz et al., 2013). Only *Salmonella* and *Campylobacter* genomes with complete allelic profiles (i.e. no truncated or missing loci) and representing unique allelic profiles were included in the final analyses (n=32,864 and n=4,134, respectively). Allelic profiles were clustered using globally optimal eBURST (goeBURST) (Francisco et al., 2009) in PHYLOViZ (Francisco et al., 2012). A distance threshold (*T*) for cluster definition, expressed as the number of allelic differences for which isolates would be considered as part of the same cluster, was applied and used to generate goeBURST clusters at all possible similarity thresholds. A “normalized” or feature-scaled *T*, expressed as a proportion of the total number of typing loci, was also calculated to facilitate comparison across datasets since each organism had a different number of loci included in its schema.

### *Campylobacter* cgMLST assay design

To identify core genes, we sampled a wide cross-section of publicly available genomes (n=5,693). Information on the strains used for core genome definition and cgMLST scheme creation can be downloaded from https://github.com/INNUENDOCON/nAWC. Draft genome assemblies were annotated using Prokka (Seemann, 2014). We used Roary (Page et al., 2015) to define a pangenome and to identify a “strict” set of core genes found in at least 99.9% of genomes in the dataset. This strict core gene definition circumvented allele assignment problems that arise when analyzing large numbers of draft genome assemblies for accessory loci that may be considered as core under a less stringent definition. A set of 697 core genes was identified based on this analysis and used to develop a cgMLST scheme for the MIST *in silico* typing platform (Kruczkiewicz et al., 2013).

### Analysis of Cluster Stability

Analysis of cluster stability was performed by using the Neighbourhood Adjusted Wallace Coefficient (nAWC), a statistic proposed here in which the Adjusted Wallace Coefficient (AWC) (Severiano et al., 2011) is used to assess cluster concordance between adjacent distance thresholds for cluster definition *n* and *n*+1, where *n* is the cut-off for number of allelic differences allowed for cluster inclusion and the nAWC is defined by:

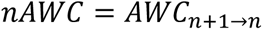

To assess cluster stability for all possible thresholds, clusters can be generated from the same matrix of allelic profiles *X* by varying the distance threshold *T^n^:*

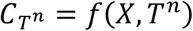

Where *f* is a function that generates *C_T^n^_* clusters from *X* at *T^n^*. The cluster concordance of clusters defined at adjacent thresholds (i.e. *T^n^* vs. *T*^*n*+1^) can be assessed for all possible distance thresholds:

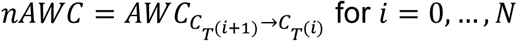

Where *N* is the maximum possible cut-off for number of allelic differences. Given *nAWC*, we can find values of *T^n^* that produce high cluster concordance with respect to *T*^*n*+1^ or, in other words, thresholds that produce stable clusters upon a shift from *T^n^* to *T*^*n*+1^.

The Shannon Index (SI) (Shannon, 1948), a measure of entropy which can be used to measure subtype diversity, was also used to examine changes in cluster partitioning as a function of the distance threshold. As a final measure of stability, at each threshold we computed the single-linkage intercluster distance for each cluster (i.e. the minimum distance to the nearest cluster) using the ‘Cluster Validation Techniques’ R package *clv* (https://CRAN.R-proiect.org/package=clv) and used these data to identify all clusters that would merge upon an increase to the adjacent threshold. For each “Merged” (*M*) cluster observed at *T*^*n*+1^ we identified: its “Recipient” (*R*) sub-cluster, i.e. the largest of its component sub-clusters observed at *T^n^*; and its “Donor” (*D*) sub-cluster(s), i.e. all remaining sub-clusters observed at *T^n^* (please refer to Fig. 3a). For each threshold, we then computed the number and size of each *M* cluster and their respective *R* and *D* sub-clusters. To examine the impact *D* sub-clusters on the nAWC at a threshold of *T*^*n*+1^, we added the total number of subtypes in *D* sub-clusters at a threshold *T^n^* and divided this value by the total number of individual subtypes in the dataset to compute the normalized aggregate size of *D* sub-clusters. A similar approach was used to compute the normalized aggregate size of *R* sub-clusters. Significance between the mean of normalized aggregate *D* sub-cluster sizes for thresholds with nAWC > 0.99 and thresholds with nAWC < 0.90 was calculated using the Mann-Whitney U test in R. The nAWC, SI, and M, *R*, *D* cluster composition statistics were computed using scripts written in the R language for statistical computing (R Core Team, 2012), available for download at https://github.com/INNUENDOCON/nAWC.

## Results

### Rapid consolidation of goeBURST clusters at low distance thresholds for cluster definition

In this study, 91,295 *Salmonella* and 6,784 *Campylobacter* genomes were analyzed using cgMLST schemas based on 330 and 697 loci, yielding 27,833 and 4,134 unique allelic profiles, respectively. These were clustered using goeBURST and cluster size and composition at all possible *T* values were then examined. As expected, a *T^0^* produced a number of singleton clusters equivalent to the number of unique profiles for *Salmonella* (n=27,833; Fig. 1a, top panel) and *Campylobacter* (n=4,134; Fig. 2a, top panel), respectively. For each organism we then observed two phases reflecting different cluster consolidation dynamics. In the first (Phase I), an increase in the threshold from *T*^0^ to *T*^1^ and to subsequent thresholds led to a steep decline in the number of clusters observed (Fig. 1a and Fig. 2a, top panel). A similar decline in Shannon Index (SI), a measure of cluster entropy, was observed (Fig. 1a and Fig. 2a, middle panel). Although a similar overall trend was observed for *Salmonella* and *Campylobacter*, the initial rate of cluster consolidation, i.e. the joining of clusters upon an increase in *T*, was much higher for the former, with a ten-fold reduction in the number of clusters (from 32,864 to 3,250) observed at a normalized *T* of 0.0667. By contrast, for *Campylobacter* a ten-fold reduction in cluster numbers (from 4,134 to 411) was observed at a normalized *T* of 0.2396. For both organisms, we then observed a secondary phase, in which cluster consolidation decreased dramatically and the number of overall clusters decreased gradually until all genomes collapsed into a single cluster. This reduction in cluster consolidation was observed at a lower normalized *T* for *Salmonella* than for *Campylobacter.* Sustained rates of cluster consolidation (i.e. below 0.1 % of total clusters) were observed by normalized thresholds of 0.1182 and 0.1679 for *Salmonella* and *Campylobacter*, respectively.

**Figure 1.**
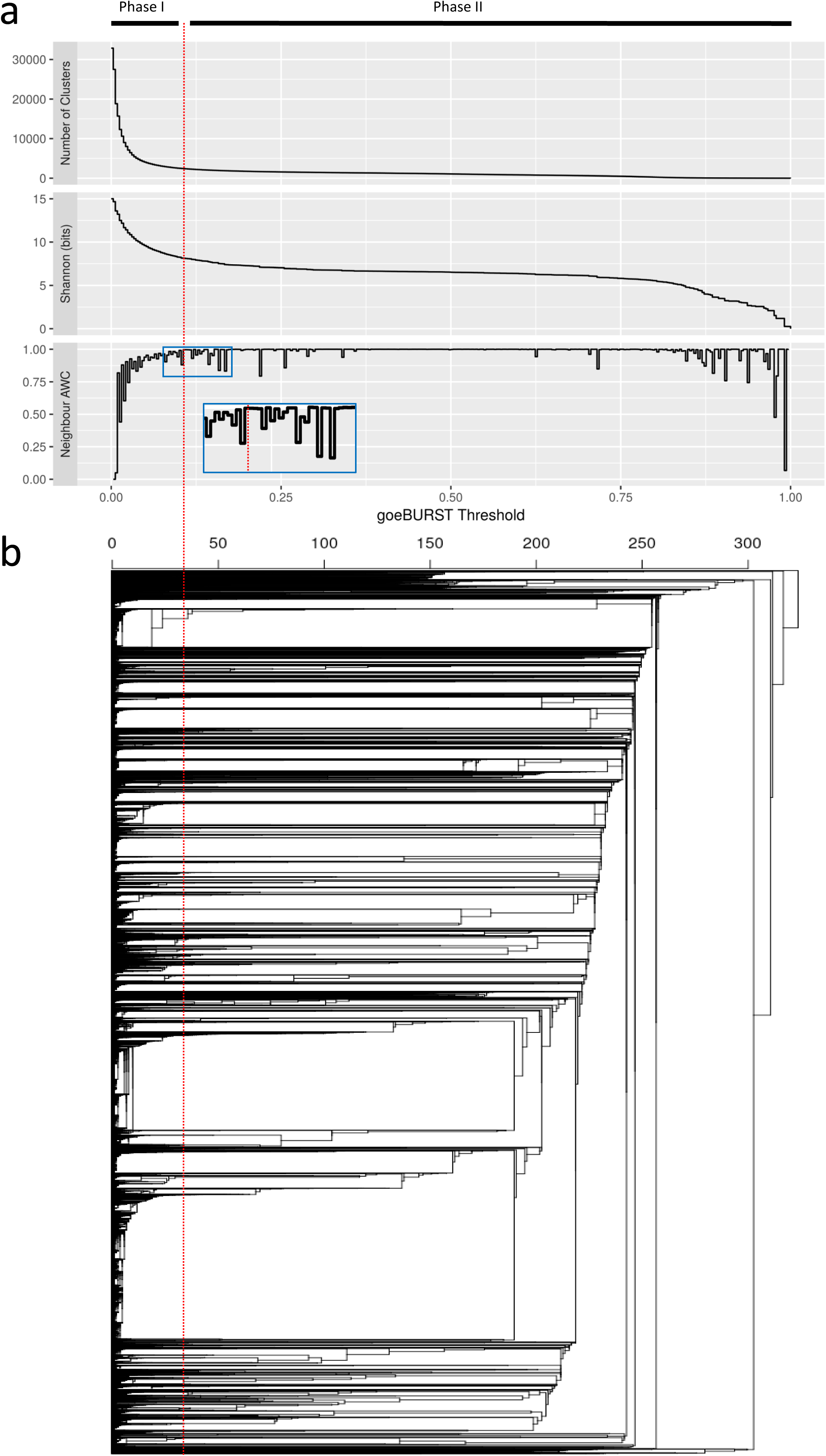
Relationship between cluster stability metrics and goeBURST threshold based on cgMLST analysis of *S. enterica.* (a) The number of clusters and Shannon Index (SI) show the impact of adjusting the distance threshold for cluster definition (*T*) on cluster stability. At the lowest *T* values, a region of high instability (Phase I) can be observed that is characterized by a rapid decline in both the number of clusters observed and the Shannon entropy. This is followed by a region of high stability (Phase II) in which cluster consolidation is greatly reduced, as evidenced by the marginal decreases in the number of clusters and Shannon entropy observed. Consistent with these cluster consolidation dynamics, Neighbourhood Adjusted Wallace Coefficient values observed in Phase I deviate from unity, as adjacent *T* values produce unstable clusters with low partition congruence. In Phase II, greater cluster stability is evidenced by sustained high nAWC values across multiple thresholds punctuated by periodic large drops in the nAWC plot (i.e. “nAWC valleys”) due to the consolidation of mid-sized clusters. The transition between Phase I and Phase II is shown (inset). (b) Cluster stability characteristics are consistent with branching patters observed in a dendrogram showing the population structure of S. *enterica.*

**Figure 2.**
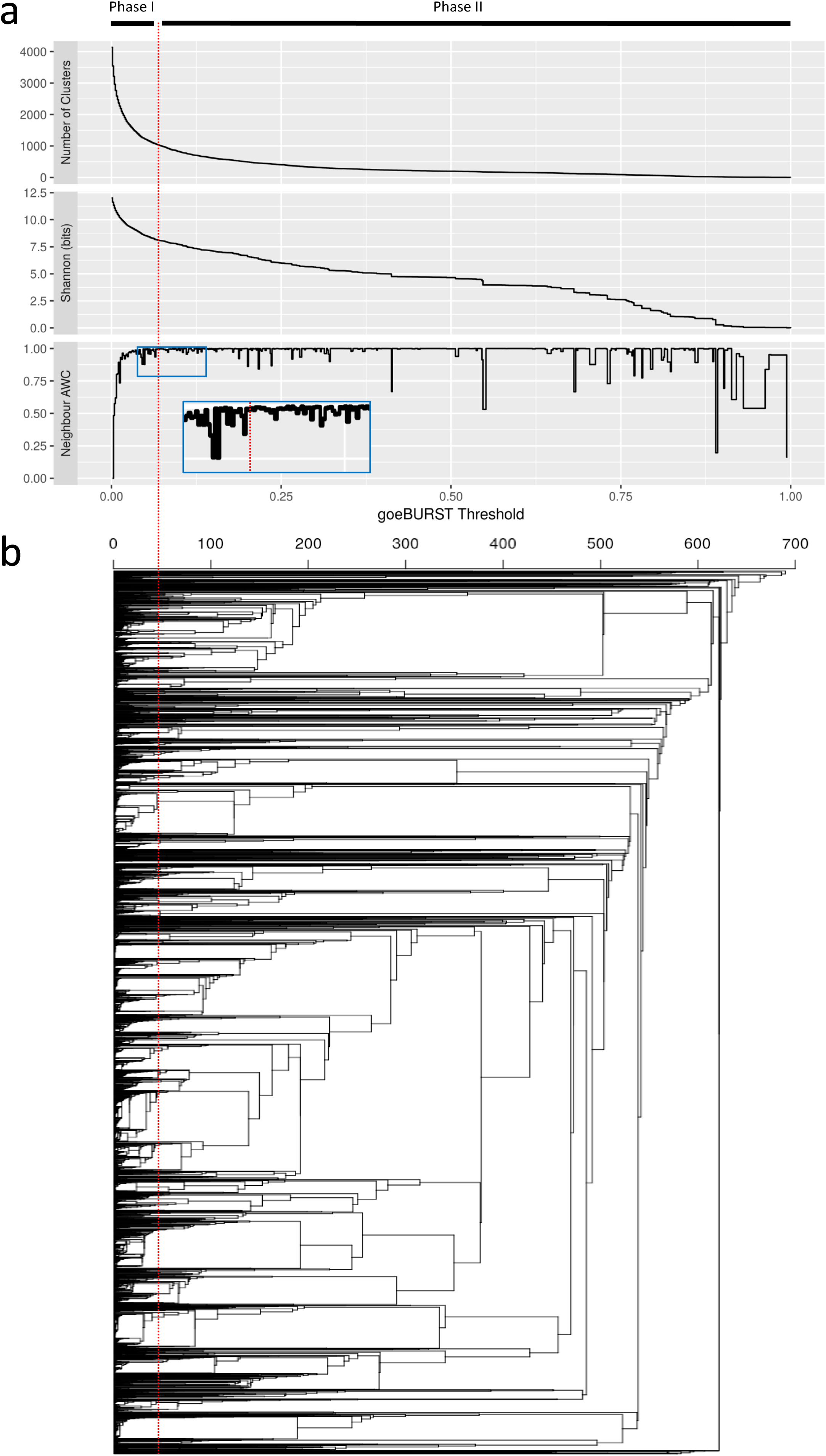
Relationship between cluster stability metrics and goeBURST threshold based on cgMLST analysis of *C. jejuni.* (a) As was observed for S. *enterica* (Fig. 1a) low distance thresholds for cluster definition (*T*) led to a region of high cluster instability (Phase I) characterized by rapid consolidation of clusters, which was followed by a region of higher stability (Phase II) in which the rate of cluster consolidation was greatly reduced. As was observed for *Salmonella*, the Neighbourhood Adjusted Wallace Coefficient (nAWC) is consistent with other measures of cluster stability, including the number of clusters observed and Shannon entropy. In Phase I, the nAWC deviated from unity while converging towards it as Phase II emerged. The transition between Phase I and Phase II is shown (inset). In Phase II, cluster stability yields sustained plateaus comprising high nAWC values with periodic “nAWC valleys”. (b) Cluster stability characteristics, which include a much larger number of nAWC valleys compared to S. *enterica* (n=44 vs. n=19, respectively), are consistent with the large number of merging lineages observed across a wider range of phylogenetic depths in a dendrogram representing the *C. jejuni* population structure.

### nAWC and SI analysis reveals threshold ranges with high levels of cluster stability

The nAWC, a statistic for measuring the level of cluster agreement between adjacent similarity thresholds, was computed for both *Salmonella* and *Campylobacter.* In the nAWC plots (Fig. 1a and Fig. 2a, bottom panels), it is possible to observe evidence consistent with the two different phases of cluster consolidation previously discussed. In Phase I, an increase from *T*^0^ to *T*^1^ and to subsequent thresholds resulted in successive nAWC values that deviated from the maximum value of 1, reflecting the low partition congruence between neighbouring thresholds consistent with high cluster instability. Values for nAWC as low as 0.3447 and 0.3285 were observed in Phase I for *Campylobacter* and *Salmonella*, respectively. As the distance threshold was increased, we then observed nAWC convergence towards unity, leading to a secondary phase defined by nAWC values that remained at or near this maximum value, consistent with highly stable cluster configurations and reduced rates of cluster consolidation. Transition to Phase II, which we defined at the earliest point at which five or more consecutive thresholds yielded nAWC values that remained above 0.9900, was observed at normalized *T* values approximating 0.0646 and 0.1061 for *Campylobacter* and *Salmonella*, respectively. In Phase II, we observed nAWC plateaus that were interrupted by isolated temporary decreases in the nAWC (i.e. nAWC “valleys”) ranging in value from 0.500 to 0.950 triggered by the merging of mid- and large-sized clusters at specific distance thresholds. Notably, we also observed large drops in Shannon Index at *T* values generating large nAWC valleys, consistent with a loss of entropy through cluster consolidation, a trend that was most conspicuous for *Campylobacter* (Fig. 2a, middle and lower panels).

### Highly asymmetrical cluster consolidation leads to nAWC values approaching unity

For each threshold we computed the minimum single-linkage intercluster distance for every cluster in order to identify all clusters expected to merge (i.e. “Merged” clusters or M) upon an increase to the adjacent threshold (i.e. *T^n^* → *T*^*n*+1^). We examined the composition of each *M* cluster with regards to the relative contribution of “Recipient” (*R*) and “Donor” (*D*) sub-clusters that comprised it (see representation in Fig. 3a). As can be seen in Fig. 3b, there is a strong negative correlation between the nAWC and the normalized aggregate size of *D* sub-clusters, i.e. the proportion of overall subtypes in *D* sub-clusters (r = −0.7103 and −0.8398 for *Campylobacter* and *Salmonella*, respectively). Consistent with this observation, the mean of the normalized aggregate size of *D* sub-clusters for thresholds with an nAWC > 0.99 was significantly smaller than that of thresholds with an nAWC < 0.90 (Fig. 3c) for both *Campylobacter* (0.0017 ± 0.0021 vs. 0.0435 ± 0.0386; p = 8.02 × 10^−21^) and *Salmonella* (0.0024 ± 0.0032 vs. 0.0969 ± 0.0802; p = 7.26 × 10^−20^).

**Fig. 3.**
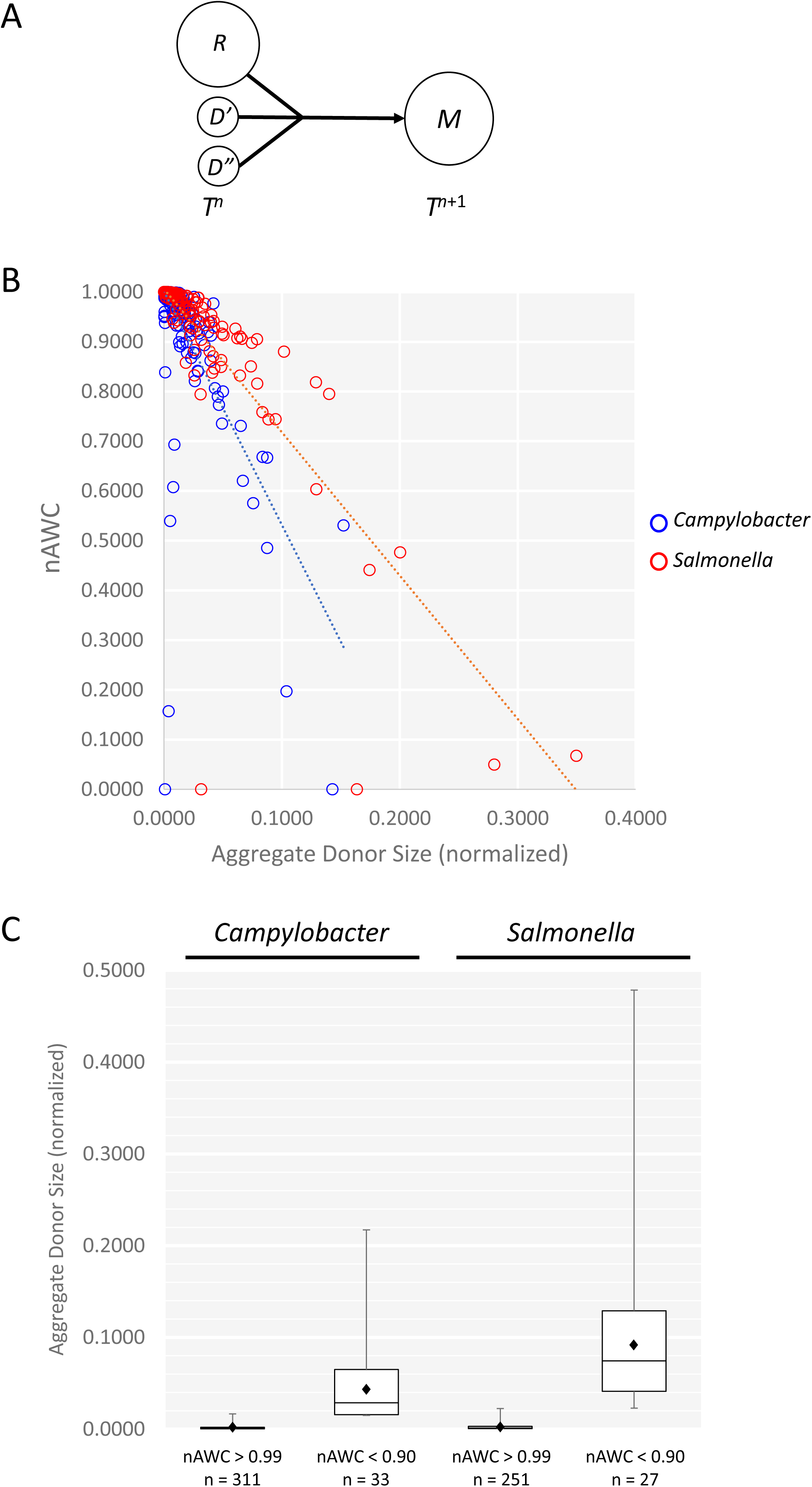
The influence of Donor cluster size on the Neighbourhood Adjusted Wallace Coefficient. (A) A “merged cluster” *M* at threshold *n*+1 results from the merging of a “recipient” cluster *R* and one (or more) “donor” sub-clusters (e.g. *D’* and *D*”) upon a shift in distance threshold from *n* (i.e. *T^n^* → *T*^*n*+1^). (B) For both *Campylobacter* and *Salmonella*, there is an inverse relationship between nAWC and the aggregate donor size (r = −0.7103 and −0.8398 for *Campylobacter* and *Salmonella*, respectively), which is expressed as the proportion of total subtypes in donor clusters contained within merged clusters to facilitate comparison between datasets. (C) A comparison of the mean aggregate donor size for thresholds with nAWC values deviating significantly from unity (i.e. nAWC < 0.9) and thresholds with nAWC values approaching unity (i.e. nAWC > 0.99) shows that the former are significantly higher for both *Campylobacter* (p = 8.02 × 10^−21^) and for *Salmonella* (p = 7.26 × 10^−20^). Boxplots represent quartile values, ♦ denotes sample mean.

### The nAWC can be used to identify distance thresholds critical to cluster consolidation

We examined *D* and *R* sub-cluster data for all *M* clusters at sets of consecutive thresholds representing isolated nAWC valleys, as described in Fig. 4a. Data for *Campylobacter* (Fig. 4b) for *M* clusters observed at thresholds of 32, 33, and 34 loci differences, which generated nAWC values of 0.9950, 0.8775, and 0.9950 respectively, show that most *M* clusters at all three thresholds resulted from the fusion of *R* sub-clusters of varying sizes to much smaller *D* sub-clusters. The primary difference between the central threshold and its flanking thresholds was a single *M* cluster comprising a *D* sub-cluster (n=48) and an *R* sub-cluster (n=164), which was sufficient to trigger a drop in the nAWC from 0.9950 to 0.8775. Similar observations could be made on the second example shown in Fig. 4c, which shows data for *Salmonella* merged clusters observed at thresholds of 33, 34, and 35 loci differences, which generated nAWC values of 0.9914, 0.8808, and 0.9966 respectively. As in the previous example, most of the merged clusters resulted from the fusion of *R* sub-clusters with small *D* sub-clusters and the main difference was at two merged clusters – (*D* (n=354) and *R* (n=391); *D* (n=631) and *R* (n=756) – that were sufficient to trigger the drop in nAWC from 0.9914 to 0.8808. Similar observations were made at other thresholds that produced nAWC valleys (results not shown).

**Fig. 4.**
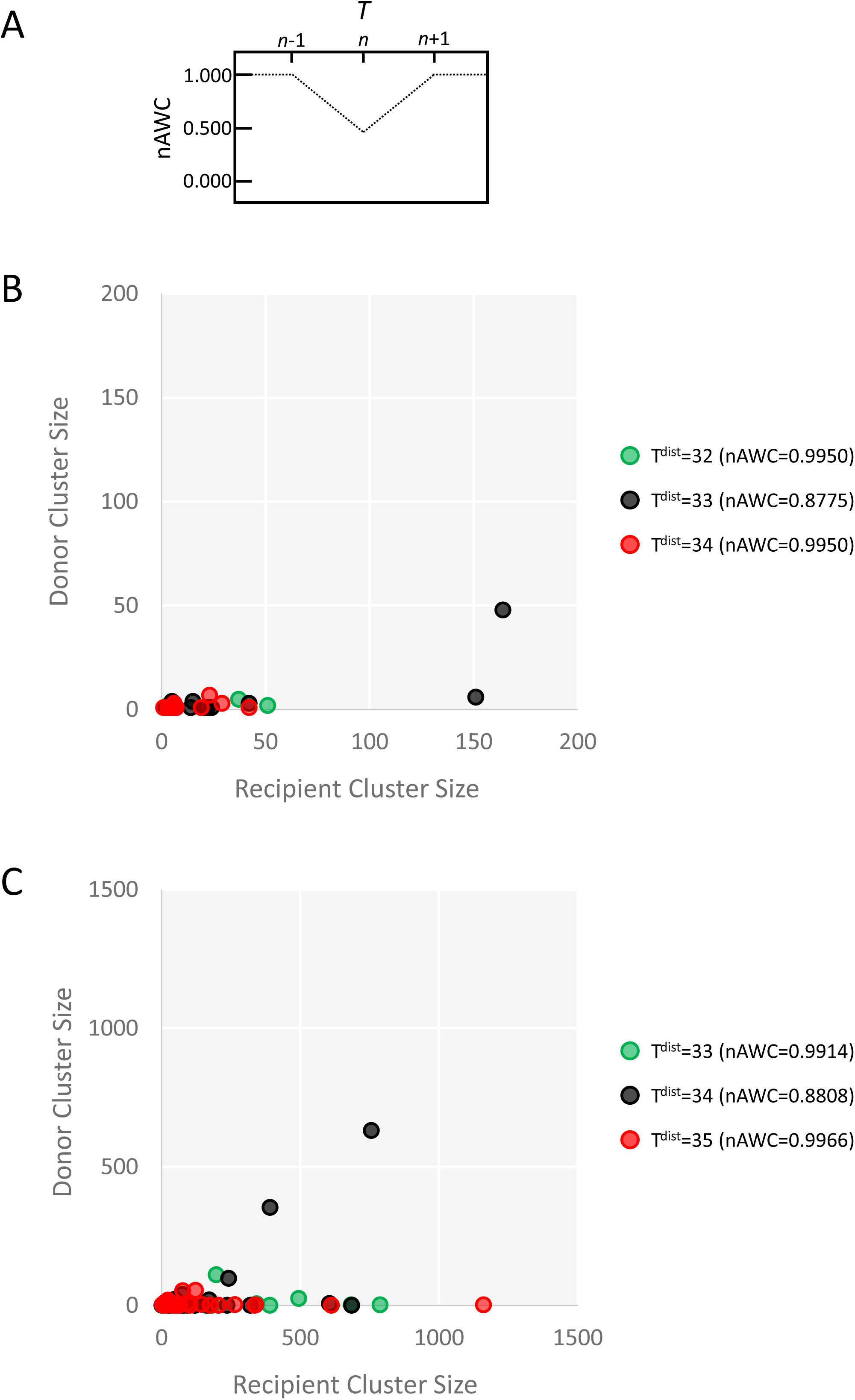
Cluster merging dynamics at critical thresholds generating large nAWC deviations. We examined cluster merging statistics (Donor size vs. Recipient size) for all merged clusters at sets of three consecutive thresholds that included a central threshold representing an “nAWC valley” flanked by thresholds with nAWC values near unity (A). For *Campylobacter* (B) and *Salmonella* (C), we observed that the majority of merged clusters represent the fusion of recipients of varying sizes with donors of much smaller size. At the threshold representing the nAWC valley, however, a small number of merged clusters with large donor sizes were sufficient to generate significant drops in the nAWC.

## Discussion

An essential part of modern infectious disease epidemiology is the establishment of stable analytical units or clusters in the population of interest that can be assigned a type name or label and that are suitable for longitudinal tracking. Once achieved, this provides a standardized means of efficiently communicating and contextualizing results between laboratories.

Subtyping methods based on whole-genome sequencing have brought a dramatic improvement over a previous generation of subtyping methods providing a more detailed view of bacterial populations and the potential for greater specificity in describing subpopulations of interest. Although the establishment of optimal thresholds for cluster definition is long-standing issue in the field of molecular epidemiology that predates WGS, WGS-based subtyping approaches exacerbate this challenge through problems of scale and resolution. Global databases of WGS data are expected to grow at a rate that is much faster than any previous historical counterpart in the pre-WGS era. At the same time, WGS-based subtyping data provide a discriminatory power that is orders of magnitude greater than legacy subtyping methods. Both the addition of novel genome sequences and adjustments in clustering thresholds for high-resolution data can have a dramatic impact on cluster composition and stability. This has the potential to influence the application of WGS-based typing data in an epidemiological context, impacting both outbreak investigations and longitudinal surveillance. Although there are a number of analytical approaches that can be used to identify lineages in a population (e.g. Cheng et al., 2013; Raj et al., 2014), in the context of epidemiological surveillance the process for identifying clusters of interest has primarily been through the application of similarity or distance thresholds for cluster inclusion/exclusion. Our goal was to develop a methodology and framework suitable for defining distance thresholds that will originate stable and reproducible nomenclatures over clusters (i.e. partitions) in a tree or graph describing subtype relationships.

Cluster stability can be thought of as the number of consecutive distance thresholds over which cluster congruence remains high. Since metrics for assessing the congruence of clusters formed by different subtyping methods already exist, we posited that adapting these well-described methods would represent an expeditious path to developing novel approaches for assessing cluster stability on large datasets. Here we describe the Neighbourhood Adjusted Wallace Coefficient (nAWC), which is based on applying the Adjusted Wallace Coefficient of Severiano et al. (Severiano et al., 2011), to examine the partition congruence between adjacent distance thresholds used for cluster definition (i.e. *T*^*n*+1^ → *T^n^*).

Using data for *Salmonella* and *Campylobacter*, we show that this approach can be used to assess cluster stability and to identify sets of distance thresholds (i.e. “neighbourhoods”) for stable cluster definition that may be suitable for longitudinal tracking of lineages of interest and that could be used in nomenclature development. In this study, 91,295 *Salmonella* and 6,784 *Campylobacter* genomes were analyzed using cgMLST, unique allelic profiles were clustered using goeBURST and clusters were generated at all possible distance thresholds for cluster definition, allowing us to study cluster stability in greater detail.

We first examined how the number of clusters changed as a function of *T* since this simple metric could be used to determine the rate of cluster consolidation (i.e. merging of clusters), a basic measure of cluster stability. For both *Salmonella* and *Campylobacter*, we observed two distinct phases (I and II) characterized by differences in the rate of cluster consolidation. In Phase I, high rates of cluster consolidation at the lowest distance thresholds led to a rapid decline in the number of clusters, as genomes with unique allelic profiles clustered into small groups of highly similar genomes and these further consolidated into fewer clusters of larger size. This region was characterized by successive thresholds with nAWC values deviating from unity but converging towards this maximum value as the rate of cluster consolidation decreased. Although this suggests that the definition of clusters in Phase I would lead to unstable nomenclatures for longitudinal tracking, the unstable clusters generated by thresholds in this region might instead be better suited for short term tracking of strains. A secondary phase (Phase II) then emerged as the datasets reached a quasi-stable cluster configuration characterized by nAWC plateaus in which successive thresholds generated nAWC values that remained at or near unity. In this phase, mid-sized clusters representing major lineages in the population increased in size primarily through merging with smaller clusters and periodically consolidating with other mid-sized clusters, thus producing concomitant drops in the nAWC, observed as “valleys” in the nAWC plot.

Overall nAWC trends observed for *Salmonella* and *Campylobacter* were very similar, but there were important distinctions that we believe are reflective of differences in their respective population structure. Although Phase II for both organisms displayed plateaus comprising nAWC values approaching unity, for *Salmonella* nAWC plateaus were longer, with fewer and smaller valleys, indicative of the much more stable subpopulations in the *Salmonella* dataset. This is consistent with the *Salmonella* population structure, which is highly clonal and characterized by lineages (i.e. eBURST groups) that are also highly concordant with serovar designations (Achtman et al., 2012; Alikhan et al., 2018). In contrast, *Campylobacter* cluster consolidation was observed across a wider range of *T* values (i.e. phylogenetic depths), as evidenced by the larger number of nAWC valleys, which are indicative of merging between mid-sized clusters, compared to *Salmonella* (n=44 vs. n=19), which is also observed in their respective dendrograms (Fig. 1b and Fig. 2b).

To further explore the effect of distance thresholds on cluster stability, we investigated cluster consolidation as a function of *T.* Consolidation between “unstable” clusters, which are expected to merge upon a shift to the adjacent distance threshold, is governed by the minimum single-linkage intercluster distance (i.e. the smallest pairwise distance between any two genomes in the respective clusters) since a *T* that exceeds this distance will trigger cluster consolidation. For each threshold *n*, we identified sets of unstable clusters, a larger “recipient” (*R*) and one or more smaller “donor” (*D*) sub-clusters producing a “merged” (*M*) cluster upon a shift to *n*+1. For both *Salmonella* and *Campylobacter*, we found that thresholds generating nAWC values deviating substantially from unity (and which thus represent significant changes in cluster partitioning) differ from thresholds with nAWC values at or near unity in fundamental ways. While, in general, the former were more likely to contain a larger number of unstable clusters, a more important determinant leading to reduced nAWC values was aggregate donor size. A small number of *M* clusters in which *D* and *R* sub-clusters were closer in size could generate an nAWC value that was discernibly reduced from the theoretical maximum; conversely, a large number of *M* clusters in which large *R* sub-clusters absorbed small *D* sub-cluster(s) had a negligible impact on the nAWC.

Thus, while an nAWC at unity implies the absence of unstable clusters, an nAWC near unity implies that unstable clusters represent a small proportion of the overall dataset and/or that cluster consolidation is primarily through the addition of *D* sub-clusters of small size relative to *R* sub-clusters. Moreover, the amplitude of the nAWC deviation from unity is dictated by the proportion of subtypes included in *D* sub-clusters in relation to the size of the overall subtype population. Our results show that cluster stability metrics can be used to identify regions of stability in which a change in the distance threshold for cluster definition will not materially affect cluster membership. Moreover, our results show the propensity for small distance thresholds (e.g. lower than 45 out of 697 alleles for *C. jejuni*, 35 out of 330 alleles for S. *enterica*) to yield clusters that are inherently unstable and likely unsuitable for long-term tracking or nomenclature development. Nonetheless, high cluster instability exhibited at small distance thresholds suggests that they can be exploited for defining clusters for highly-specific short-term tracking of strains of interest, as in the case of outbreak investigations.

Global WGS-based surveillance will require robust approaches for defining nomenclatures based on stable clusters that both harness the immense leap in discriminatory power afforded by WGS data while allowing for robust tracking of strains of interest; our results suggest that the nAWC can be used to identify thresholds that can accommodate these needs. Using datasets from *Salmonella enterica* and *Campylobacter jejuni*, we show that clusters generated using such thresholds are both stable and reflect basic units in the overall population structure of the respective species. As these represent both strongly and weakly clonal bacterial species, our results suggest that the approach could be useful for defining robust clusters that can form the basis for the stable nomenclatures that are essential to global surveillance networks for pathogen tracking.

## Acknowledgments

The Authors wish to acknowledge EnteroBase, the NCBI’s Short Read Archive and their contributors, without whom this work would not be possible. The authors wish to acknowledge CSC – IT Center for Science, Finland, for computational resources. Financial support for this work was provided through the Government of Canada’s Genomics Research and Development Initiative (ENT, DORB, PK). FP has been supported by the University of Bologna PhD Programme grant. JAC and MR have been supported by INNUENDO project co-funded by the European Food Safety Authority (EFSA), grant agreement GP/EFSA/AFSCO/2015/01/CT2 (“New approaches in identifying and characterizing microbial and chemical hazards”). The conclusions, findings, and opinions expressed in this review paper reflect only the view of the authors and not the official position of the European Food Safety Authority (EFSA). JAC was partially supported by the following projects: ONEIDA project (LISBOA-01-0145-FEDER-016417) co-funded by FEEI - “Fundos Europeus Estruturais e de Investimento” from “Programa Operacional Regional Lisboa 2020” and by national funds from FCT - “Fundação para a Ciência e a Tecnologia” and BacGenTrack (TUBITAK/0004/2014) [FCT/ Scientific and Technological Research Council of Turkey (Türkiye Bilimsel ve Teknolojik Aragrrma Kurumu, TÜBITAK)].

The authors have no conflicts of interest to declare.

